# Autistic children show reduced activity in left superior temporal cortex in response to social-emotional meaning in language

**DOI:** 10.64898/2026.01.02.697285

**Authors:** Hanlin Wu, Matthew Ganquan Shi, Phoebe Wai-Yu Chan, Peilun Song, Siu-Fung Lau, Gary Yu-Hin Lam, Xin Kang, Patrick CM Wong, Xiujuan Geng

**Affiliations:** Department of Linguistics and Modern Languages, The Chinese University of Hong Kong; Brain and Mind Institute, The Chinese University of Hong Kong; Department of Linguistics, The University of Maryland, College Park; Department of Educational Psychology, The Chinese University of Hong Kong; Research Centre for Language, Cognition, and Language Application, Chongqing University; Key Laboratory of Hearing, Speech, and Cognition of Chongqing Municipal Health Commission, Chongqing, China

**Keywords:** Autism, Children, Social-emotional processing, Language comprehension, Superior temporal sulcus

## Abstract

Autistic children often differ from non-autistic peers in social interaction, emotional processing, and language communication. While the neural differences between autistic and non-autistic children in processing paralinguistic social-emotional cues, such as emotional prosody, are well-documented, much less is known about how the brain processes the social-emotional meaning carried by the language content itself. In non-autistic adults, the left superior temporal sulcus (STS) and adjacent gyrus show a social-emotional bias, responding more strongly to social-emotional content than to content describing inanimate objects. We investigated whether this bias is present in children and whether it is altered in autistic children. We recruited 78 right-handed children who underwent functional magnetic resonance imaging (fMRI) scanning while listening to sentences describing either people in emotional situations (social-emotional sentences) or non-human objects, events, or concepts (object sentences). After excluding those with excessive head motion, we analyzed data from 58 of them (37 autistic and 21 non-autistic; mean age 10 years). Across the sample as a whole, the left STS responded more strongly to social-emotional sentences than to object sentences, indicating that this social-emotional bias is already present in childhood. Importantly, this bias was reduced in autistic children compared to non-autistic controls, whereas no group difference was observed in general auditory or lexico-syntactic processing. This reduced activity in the left STS in response to social-emotional semantics may contribute to autistic children’s social communication challenges.

## INTRODUCTION

Autistic children often differ from non-autistic peers in social interaction and joint attention (McEvoy et al., 1993; Mundy et al., 1986; Travis & Sigman, 1998), in emotional processing (Corbett et al., 2009; Kuusikko et al., 2009), and in language communication (Loveland & Landry, 1986; Schaeffer et al., 2023). To understand the mechanisms underlying these challenges, research has been approaching them from two directions. On the one hand, psycholinguistic research has focused on the structural components of language, such as phonology, morphology, and syntax, revealing heterogeneous language profiles among autistic individuals (Boucher, 2012). Autistic children with grammatical impairment did not show greater autism symptom levels than those without it (Wittke et al., 2017). On the other hand, research on social cognition has extensively examined how autistic individuals process non-linguistic or paralinguistic social cues, identifying atypical recognition of facial emotions (Harms et al., 2010; Lozier et al., 2014) and atypical processing of emotional prosody and vocal affect (Matsumoto et al., 2016; McCann et al., 2007; Portnova et al., 2023). However, situated between these two well-studied domains lies an important yet frequently overlooked intersection: the processing of the social and emotional meaning conveyed by the language content itself.

Real-world communication relies heavily on the ability to understand words and sentences that describe human relationships (e.g., “friend”), interactions (e.g., “hug”) and emotional states (e.g., “grief”, “joy”). The processing of such social-emotional semantics requires more than representing the linguistic information; it requires representing the social and emotional meaning that the words and sentences convey. In language comprehension, this social and emotional significance is not a separate add-on to lexical meaning but an integral part of it: emotional content is part of the semantic representation of abstract words (Kousta et al., 2011; Ponari et al., 2018) and modulates word recognition from early, largely automatic stages (Citron, 2012).

However, evidence suggests this processing is functionally distinctive in autistic individuals. A systematic review of emotional language processing (Lartseva et al., 2015) found that explicit valence classification was preserved in autism in about half the studies, whereas the automatic advantages were more consistently reduced. In an event-related potential study (Yeh et al., 2026), autistic and non-autistic children and adolescents judged the valence of written emotional words. Group differences were trend-level, but across the sample the late positivity for positive emotion-label words tracked social difficulties, which was read as a selective difficulty in understanding emotional concepts from language. This pattern suggests that while the semantic information of these words is retrieved successfully, the subsequent cognitive evaluation of their emotional significance might be altered.

The neural architecture supporting this integration of language and social affect has been associated with the left superior temporal gyrus (STG) and superior temporal sulcus (STS). The STG and STS are engaged bilaterally in the acoustic and phonological analysis of speech. Acoustic-phonetic features are encoded in the bilateral STG (Bhaya-Grossman & Chang, 2022), and phonological processing has been proposed for the bilateral mid-posterior STS (Hickok & Poeppel, 2007). At the sentence level, the left anterior STS responded more strongly to sentences than to word lists (Humphries et al., 2006), and left anterior temporal cortex responded more strongly to normal sentences than to scrambled sentences (Vandenberghe et al., 2002).

Anterior temporal regions have also been suggested in representing social meaning. Zahn et al. (2007) localized social-concept knowledge to the superior anterior temporal cortex, and Rice et al. (2018) localized person knowledge to ventral anterior temporal cortex, arguing for graded specialization across a transmodal hub rather than a discrete social module. Superior temporal activity has also been found to decrease as autism level increases (Kausel et al., 2024; Zilbovicius et al., 2006).

To disentangle linguistic processing from the processing of social-emotional meaning, Mellem et al. (2016) had non-autistic adults silently read stimuli that varied in constituent size (from unstructured word lists to full sentences) and in content type (social-emotional, social, object-related, or jabberwocky). The left STG and STS showed a social-emotional bias, responding significantly more to content describing people in emotional situations than to content describing inanimate objects.

However, it remains unknown whether this social-emotional bias in the left STG/STS is already present during childhood and whether it is altered among autistic children. We hypothesize that this social-emotional bias may already be present in childhood as previously observed in non-autistic adults. We further hypothesize that this bias may be reduced in autistic children relative to their non-autistic peers.

The current study aims to test these hypotheses by investigating the neural processing of social-emotional versus object sentences in Cantonese-speaking autistic and non-autistic children. We used functional Magnetic Resonance Imaging (fMRI) and speech stimuli that controlled for syntactic complexity and lexical frequency to isolate the neural response to semantic content. We predict that while both autistic and non-autistic children may demonstrate the expected social-emotional bias in the left STG/STS like in non-autistic adults, autistic children may particularly show reduced activity in this region for social-emotional content compared to non-autistic peers.

## METHODS

### Participants

We recruited 78 right-handed children (age range 7.97 to 12.47 years, mean age 10.01 years), of whom 54 had previously received an autism diagnosis (49 male, 5 female) and 24 served as non-autistic controls (19 male, 5 female). All participants were native Cantonese speakers with hearing within normal range and were recruited from mainstream schools in Hong Kong. Written informed consent was obtained from a parent or guardian of each participant prior to any experiments. The study protocol adhered to the ethical standards of the Helsinki Declaration and was approved by the Joint Chinese University of Hong Kong-New Territories East Cluster Clinical Research Ethics Committee.

### Behavioral Assessments

Qualified therapists administered assessments of autism symptom level and linguistic ability. Autism symptom level was assessed with the Autism Diagnostic Observation Schedule Second Edition, Module 3 (ADOS-2) (Lord et al., 2012). Linguistic ability was assessed with the Hong Kong Cantonese Oral Language Assessment Scale, Textual Comprehension subscale (HKCOLAS) (T’sou et al., 2006). Additionally, their non-verbal IQ was assessed with the Test of Nonverbal Intelligence-Fourth Edition (TONI-4) (Brown et al., 2010). In addition, a parent-report language background questionnaire was used to rate each child’s proficiency in Cantonese and English on listening, speaking, reading, and writing, each on a scale from 0 to 7. For each language we examined the four dimensions individually and a composite score (mean score across the four dimensions), together with a Cantonese-minus-English dominance index (Table S3).

### Experimental Design and Materials

While in the MRI scanner, all participants listened to speech stimuli as fMRI scans were acquired. The stimuli consisted of two types (see Table 1 for examples; see Table S1 for the full list): (a) *social-emotional sentences* that described people in emotional situations (half positive, half negative); and (b) *object sentences* that described non-human objects, events, or concepts. Each type comprised 36 oral Cantonese sentences, each exactly eight characters long (equivalent to eight syllables). The two sentence types were matched for mean word frequency (Lai & Winterstein, 2020) (*t*(70) = 0.22, *p* = 0.824), as a measure of general lexical-semantic processing difficulty, and mean dependency distance (Liu, 2008) (*t*(70) = 1.12, *p* = 0.266), as a measure of general syntactic complexity.

**Table 1.** Examples of experimental materials (English translations)

| Type | Example (English translation) |
| --- | --- |
| Social-emotional sentences |  |
| Positive | The doctor received many compliments.<br>My older sister laughed at the joke.<br>Ka-Kei was thrilled by her success. |
| Negative | The driver crashed the car in haste.<br>Mom has so many worries.<br>Chi-Keung feels he is no good. |
| Object sentences |  |
|  | There is a ball inside the square box.<br>The Earth revolves around the Sun.<br>Banana peels are yellow.<br>Paper is made from wood.<br>The sun rises in the east.<br>Tree roots lie buried beneath the soil. |

All sentences were synthesized with Microsoft Azure Text-to-Speech using a single female Cantonese voice, a method that has been used in previous studies of spoken-language comprehension for audio control (e.g., Wu et al., 2024, 2025; Wu & Cai, 2026). Speaker identity, voice quality, and prosody were therefore consistent across all items and conditions, so the two sentence types differed in semantic content but not in emotional prosody or other paralinguistic cues. All stimuli were checked by a native Hong Kong Cantonese-speaking author (P.W.Y. Chan) and confirmed to be free of perceptible unnaturalness. Additionally, reversed speech was included as a non-linguistic auditory control, created by temporally reversing each of the 36 object sentences.

The experiment employed a block design. The task was 8.4-minute long and included 6 cycles (1.4 min each). Each cycle began with a 12-s silence block, followed by the three main 24-s experimental blocks: a social-emotional sentence block, an object sentence block, and a reversed speech block. The presentation order of these three experimental blocks was counterbalanced across the six cycles and across all participants.

The social-emotional sentence, object sentence, and reversed speech blocks each contained six trials. A single trial consisted of a 3500 ms presentation period followed by a 500 ms blank screen. At the onset of the presentation period, the auditory stimulus began playing, and a fixation cross appeared on the screen. While the auditory stimulus was shorter (mean duration = 2180 ms), the fixation cross remained visible for the full 3500 ms duration. Throughout all trials, participants were instructed to listen attentively to the auditory stimuli without making behavioral response or judgment during scanning. The silence blocks served as a baseline and contained three trials where the fixation cross was displayed for 3500 ms in silence, each followed by a 500 ms blank screen.

### MRI Acquisition

MR images were collected on a 3 T Siemens Magnetom Prisma scanner at the Prince of Wales Hospital, Hong Kong. T1-weighted images (T1w) with MPRAGE and functional images with EPI BOLD were collected. Main imaging parameters of T1w were as follows: repetition time (TR) = 1900 ms; echo time (TE) = 2.38 ms; flip angle = 9°; field of view (FOV) = 230 × 230 × 172 mm; voxel size = 0.9 × 0.9 × 0.9 mm; acquisition matrix = 256 × 256 × 192. The functional MRI parameters were as follows: TR = 1000 ms; TE = 83 ms; flip angle = 61°; FOV = 220 × 220 × 140 mm; voxel size = 2 × 2 × 2 mm; acquisition matrix = 110 × 110 × 70; simultaneous multi-slice acquisition with a multiband factor of 5.

### Head Motion Assessment and Data Exclusion

All functional images underwent realignment to the mean image using six-parameter rigid-body transformation (3 translations and 3 rotations). Frame-wise displacement (FD) was calculated as the sum of absolute differences in the six movement parameters between consecutive volumes, with rotational parameters converted to millimeter displacement assuming a 50 mm head radius (Power et al., 2012). Participants were excluded from further analysis based on the following motion criteria: (a) mean FD exceeding 0.5 mm across the entire scan session; (b) maximum FD exceeding 5 mm for any single volume-to-volume transition; or (c) more than 25% of volumes exceeding an FD threshold of 0.5 mm. Of the 78 children recruited, 20 were excluded due to excessive head motion, resulting in a final sample of 58 participants (37 autistic children and 21 non-autistic controls). Motion-related exclusion was more frequent in the autistic group (17 of 54, 31%) than in the non-autistic group (3 of 24, 13%). Of this final sample, non-verbal IQ (TONI-4) was available for 57 participants (36 autistic, 21 non-autistic) and missing for one autistic child; HKCOLAS scores were available for 50 participants (29 autistic, 21 non-autistic) and missing for eight autistic children, and the child without a TONI-4 score was among these eight. All 58 participants completed the ADOS-2. Co-occurring conditions were surveyed but not exclusionary: 27 of 37 autistic children (73%) had at least one, most commonly ADHD, whereas no non-autistic child had any co-occurring diagnosis (Table S7).

### MRI Preprocessing

T1-weighted and functional images were preprocessed using SPM12 (Ashburner et al., 2014) in MATLAB. Following realignment and motion assessment, functional images were corrected for slice timing. Each participant’s T1-weighted image was coregistered to their mean functional image and then segmented. The deformation field estimated during segmentation was used to normalize the functional images to Montreal Neurological Institute (MNI) standard space, using the adult SPM12 tissue probability map template with East-Asian affine regularization. These normalized images were then spatially smoothed with a 6 mm full-width at half-maximum (FWHM) Gaussian kernel.

### Whole Brain Analyses

#### First-level Analysis

Individual participant data were analyzed using a general linear model (GLM) implemented in SPM12 (Ashburner et al., 2014). The GLM included four experimental conditions: Silence (Sil, baseline); Reversed speech (Rev, non-linguistic auditory control); Object sentences (Obj); and Social-emotional sentences (SE). Each condition was modeled with boxcar waveforms convolved with the canonical hemodynamic response function. Events were modeled with boxcars of each stimulus’s actual duration (1.98 to 2.36 s; silence 3.5 s). Three linguistic variables (z-scored mean word frequency, mean dependency distance, and word count) for each sentence were included as nuisance regressors to control for potential confound from lexical and syntactic processing. Additionally, six rigid-head motion parameters, FD values, and the binary FD flags (volumes with FD > 0.5 mm) were included as nuisance regressors to control for motion-related confounds. A high-pass filter with a 128-s cutoff was applied to remove low-frequency signal unrelated to the task, such as scanner drift and slow physiological fluctuation. Four t contrasts were estimated for each participant and entered the second-level and ROI analyses: (a) Reversed speech > Silence, testing non-linguistic auditory processing; (b) Object sentences > Reversed speech, testing general linguistic processing; (c) Social-emotional sentences > Reversed speech, testing linguistic processing of social-emotional sentences against the same acoustic baseline; and (d) Social-emotional sentences > Object sentences, which isolates the processing of social-emotional semantics.

#### Second-level Analysis

Each contrast was analyzed at the group level with a multiple regression (general linear model). The model included Group (autistic, non-autistic) as a predictor of interest, together with Sex, Age, mean FD, and non-verbal IQ as covariates (all covariates mean-centered). One autistic participant did not complete the TONI-4; in the whole-brain models the missing non-verbal IQ was replaced with the mean of the full analyzed sample (both groups combined), whereas the region-of-interest regressions use complete cases and therefore run at N = 57, or fewer when a focal predictor was missing. A sensitivity analysis excluding this child (*N* = 57) left the results essentially unchanged (see Table S8). The across-group effect identified activity common to both groups, and the diagnostic-group predictor tested for differences between autistic and non-autistic children. Statistical maps were generated for each contrast of interest. Whole-brain clusters were formed at a voxel-level threshold of *p* < 0.001 and considered significant if they survived cluster-level family-wise error correction at *p* < 0.05. Anatomical labels for cluster peaks were assigned with the AAL3 atlas (Rolls et al., 2020) and cross-checked against the HCP-MMP1 parcellation (Glasser et al., 2016), by the rule stated in the Table 3 note. We additionally estimated each contrast of interest within each group separately, as a one-sample model including the same covariates (sex, age, mean FD, and non-verbal IQ) and the same cluster-forming and cluster-level FWE thresholds as the across-group maps. The social-emotional contrasts were additionally compared between the groups with a two-sample model at the same thresholds.

### Region of Interest (ROI) Analysis

#### ROI selection

We examined the four left superior temporal peaks that Mellem et al. (2016) reported for the main effect of content type (middle STS [m-STS], anterior STG [a-STG], anterior STS [a-STS], and posterior STS [p-STS]), whose Talairach coordinates we converted to MNI space with the BioImage Suite converter, which implements the mapping of Lacadie et al. (2008). These four regions were selected independently of the present data, and every region was tested for every contrast, so the region definitions cannot be influenced by the group comparison they are used to test (Kriegeskorte et al., 2009). For each ROI we extracted each participant’s mean contrast estimate from a 6 mm sphere centered on the coordinate, averaging over the voxels within the sphere.

#### Group and individual difference analyses

Group differences were tested with two-sample *t* tests and corrected for false discovery rate (FDR) across the four regions within each contrast. To further analyze the ROI group differences we fitted linear regression models, using one consistent set of nuisance covariates in every regression model, Age, Sex, mean FD, and non-verbal IQ, with Group modeled as a fixed effect. We treated autism symptom level (ADOS-2) and language ability (HKCOLAS) as two focal predictors of equal interest. For each focal predictor we tested (i) whether it predicted ROI activity within each group, fitted separately in the autistic and non-autistic groups; (ii) whether the group difference remained when the predictor and its interaction with group were entered; and (iii) whether it was associated with activity in the pooled sample ignoring group. We repeated the analysis with the ADOS-2 social-affect (ADOS-SA) and restricted-repetitive-behavior (ADOS-RRB) subscales in place of the total score. All continuous predictors were mean-centered.

#### Valence analysis

To test whether the group difference depended on emotional valence, we re-estimated the first-level GLM with the social-emotional condition divided into positive and negative sentences, from which the [Social-emotional positive > Object] and [Social-emotional negative > Object] contrasts were analyzed. This model used the same preprocessed images and nuisance regressors as the main model and differed only in splitting the social-emotional condition by valence. For each ROI we extracted these contrast estimates and fitted a 2 (Valence: positive, negative; within-subject) by 2 (Group: autistic, non-autistic; between-subject) mixed ANOVA, and tested the simple effect of Group within each level of Valence (FDR-corrected across the four regions within each valence).

#### Smoothing sensitivity analysis

Because spatial smoothing could in principle carry signal from neighboring cortex into a region of interest, we repeated the ROI analysis on unsmoothed data: each participant’s first-level model, including the valence-split model, was re-estimated on the normalized but unsmoothed images with an otherwise identical design (Table S9).

## RESULTS

### Demographics and Behavioral Assessments

Among the 58 participants included in the final analyses (37 autistic, 21 non-autistic), the two groups did not differ significantly in age or sex distribution, but the autistic group had lower non-verbal IQ scores (*p* = 0.047; Table 2), and the non-verbal IQ was included as a covariate in every whole-brain model and in every regression model. As expected, autistic participants showed significantly higher ADOS-2 scores than non-autistic controls for total, social affect (SA), and restricted repetitive behavior (RRB) scores. Autistic participants also scored significantly lower on the HKCOLAS textual comprehension measure (Table 2). The autistic group was rated lower on five of twelve parent-reported proficiency dimensions at an uncorrected threshold, including the English and Cantonese composites, although no dimension survived false-discovery-rate correction across the twelve tests (minimum corrected *p* = 0.09). Cantonese-over-English dominance did not differ between groups (Table S3).

**Table 2.** Demographics and behavioral assessments of autistic and non-autistic children.

| Variable | Autistic | Non-autistic | *t* | *df* | *p* |
| --- | --- | --- | --- | --- | --- |
|  | Mean (*SD*) | Mean (*SD*) |  |  |  |
| Age | 10.32 (1.36)<br>[8.25, 12.47] | 10.07 (1.22)<br>[8.16, 12.28] | 0.72 | 45.62 | 0.478 |
| Sex, male / female | 32 (86%) / 5 (14%) | 16 (76%) / 5 (24%) | - | - | 0.471 |
| Non-verbal IQ | 114.89 (13.25) | 121.10 (9.60) | -2.04 | 52.27 | 0.047 |
| ADOS-2 Total | 12.38 (3.31) | 1.90 (1.55) | 16.35 | 54.50 | < 0.001 |
| ADOS-2 SA | 10.05 (3.27) | 1.38 (1.50) | 13.77 | 54.20 | < 0.001 |
| ADOS-2 RRB | 2.32 (1.43) | 0.52 (0.75) | 6.27 | 55.75 | < 0.001 |
| HKCOLAS | 21.48 (5.90) | 27.43 (3.85) | -4.30 | 47.59 | < 0.001 |
| Mean FD (mm) | 0.252 (0.075) | 0.250 (0.073) | 0.10 | 42.59 | 0.924 |
| High-motion volumes (%) | 6.65 (5.69) | 7.08 (6.47) | -0.25 | 37.39 | 0.805 |

*Group comparisons are Welch (unequal-variance) *t* tests; sex was compared with Fisher's exact test. Values in square brackets are the range. A positive *t* indicates a higher value in the autistic group. All measures cover the full sample (37 autistic, 21 non-autistic) except non-verbal IQ (36 and 21) and HKCOLAS (29 and 21). Non-verbal IQ was assessed with the Test of Nonverbal Intelligence, Fourth Edition (TONI-4). Mean FD, mean framewise displacement; high-motion volumes, percentage of the functional volumes with framewise displacement > 0.5 mm; ADOS-2, Autism Diagnostic Observation Schedule, Second Edition (Module 3); SA, social affect; RRB, restricted and repetitive behavior; HKCOLAS, Hong Kong Cantonese Oral Language Assessment Scale (Textual Comprehension).*

### Whole-Brain Effects of Sentence Type and Group

Across autistic and non-autistic groups, reversed speech elicited significant activity in bilateral auditory cortex along the superior temporal gyri (STG, Figure 1A). Compared to reversed speech, object sentences elicited heightened activity in one large left-lateralized cluster centered on the middle temporal gyrus (MTG) and extending into the superior temporal, inferior frontal, precentral, and inferior temporal cortex, together with separate clusters in the left supplementary motor area (SMA), the right fusiform gyrus, the right cerebellum, and the right postcentral gyrus (Figure 1B). Social-emotional sentences, compared to reversed speech, activated the left STS along its middle and posterior extent, with smaller clusters in the right anterior STS and the right cerebellum (Figure 1C). Contrasting the two sentence types, a single left superior temporal cluster, with its peak in the m-STS and additional local maxima extending anteriorly along the STS (−52, −6, −16) and into the superior temporal pole (−50, 10, −20), showed heightened activity for social-emotional over object sentences (Figure 1D, Figure 3A; Table 3).

**Table 3.**
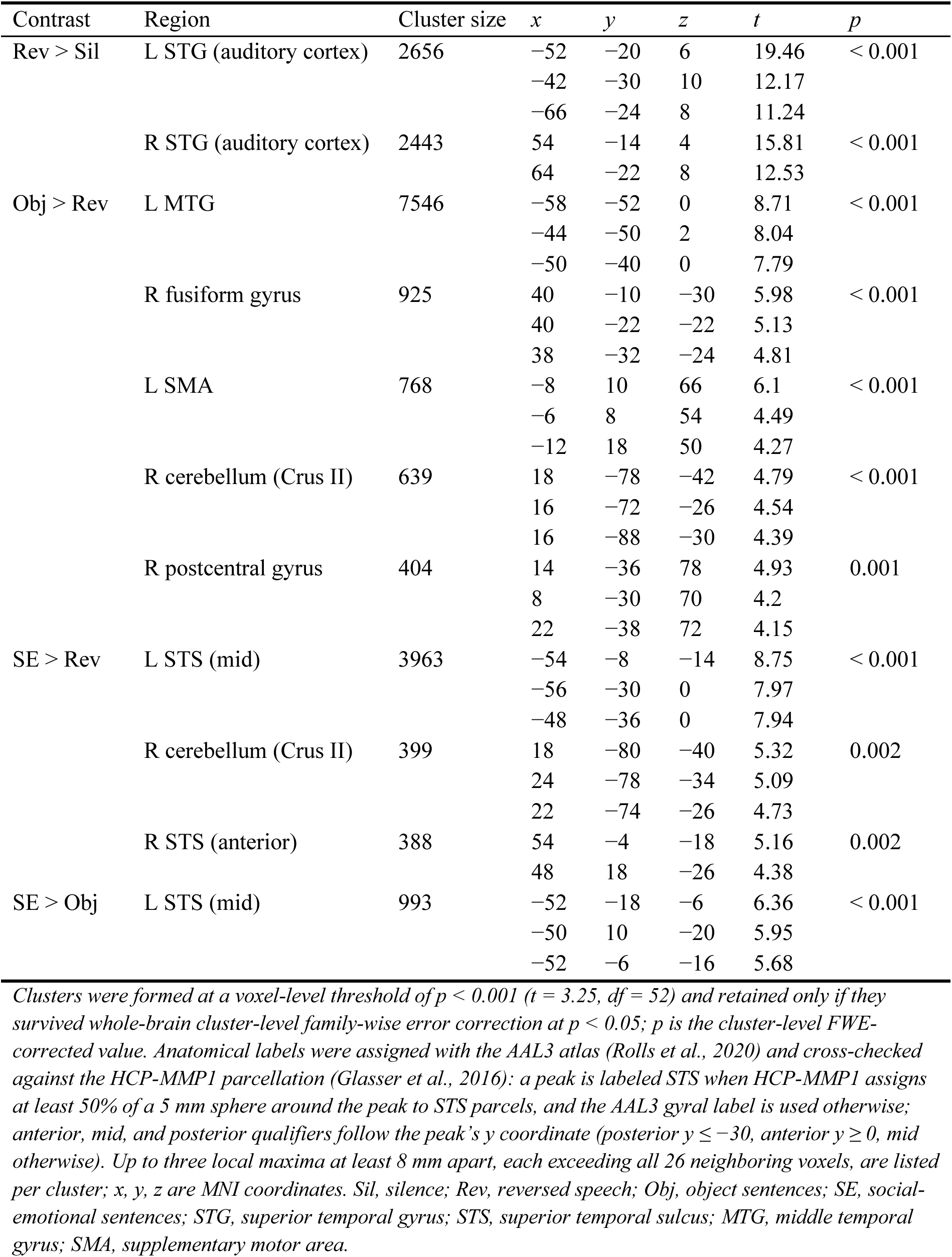
Clusters and local maxima with significant activity in each contrast across all participants (*N* = 58).

| Contrast | Region | Cluster size | $x$ | $y$ | $z$ | $t$ | $p$ |
| --- | --- | --- | --- | --- | --- | --- | --- |
| Rev > Sil | L STG (auditory cortex) | 2656 | -52 | -20 | 6 | 19.46 | < 0.001 |
|  |  |  | -42 | -30 | 10 | 12.17 |  |
|  |  |  | -66 | -24 | 8 | 11.24 |  |
|  | R STG (auditory cortex) | 2443 | 54 | -14 | 4 | 15.81 | < 0.001 |
|  |  |  | 64 | -22 | 8 | 12.53 |  |
| Obj > Rev | L MTG | 7546 | -58 | -52 | 0 | 8.71 | < 0.001 |
|  |  |  | -44 | -50 | 2 | 8.04 |  |
|  |  |  | -50 | -40 | 0 | 7.79 |  |
|  | R fusiform gyrus | 925 | 40 | -10 | -30 | 5.98 | < 0.001 |
|  |  |  | 40 | -22 | -22 | 5.13 |  |
|  |  |  | 38 | -32 | -24 | 4.81 |  |
|  | L SMA | 768 | -8 | 10 | 66 | 6.1 | < 0.001 |
|  |  |  | -6 | 8 | 54 | 4.49 |  |
|  |  |  | -12 | 18 | 50 | 4.27 |  |
|  | R cerebellum (Crus II) | 639 | 18 | -78 | -42 | 4.79 | < 0.001 |
|  |  |  | 16 | -72 | -26 | 4.54 |  |
|  |  |  | 16 | -88 | -30 | 4.39 |  |
|  | R postcentral gyrus | 404 | 14 | -36 | 78 | 4.93 | 0.001 |
|  |  |  | 8 | -30 | 70 | 4.2 |  |
|  |  |  | 22 | -38 | 72 | 4.15 |  |
| SE > Rev | L STS (mid) | 3963 | -54 | -8 | -14 | 8.75 | < 0.001 |
|  |  |  | -56 | -30 | 0 | 7.97 |  |
|  |  |  | -48 | -36 | 0 | 7.94 |  |
|  | R cerebellum (Crus II) | 399 | 18 | -80 | -40 | 5.32 | 0.002 |
|  |  |  | 24 | -78 | -34 | 5.09 |  |
|  |  |  | 22 | -74 | -26 | 4.73 |  |
|  | R STS (anterior) | 388 | 54 | -4 | -18 | 5.16 | 0.002 |
|  |  |  | 48 | 18 | -26 | 4.38 |  |
| SE > Obj | L STS (mid) | 993 | -52 | -18 | -6 | 6.36 | < 0.001 |
|  |  |  | -50 | 10 | -20 | 5.95 |  |
|  |  |  | -52 | -6 | -16 | 5.68 |  |

Clusters were formed at a voxel-level threshold of $p < 0.001$ ( $t = 3.25$ , $df = 52$ ) and retained only if they survived whole-brain cluster-level family-wise error correction at $p < 0.05$ ; $p$ is the cluster-level FWE-corrected value. Anatomical labels were assigned with the AAL3 atlas (Rolls et al., 2020) and cross-checked against the HCP-MMP1 parcellation (Glasser et al., 2016): a peak is labeled STS when HCP-MMP1 assigns at least 50% of a 5 mm sphere around the peak to STS parcels, and the AAL3 gyral label is used otherwise; anterior, mid, and posterior qualifiers follow the peak's $y$ coordinate (posterior $y \leq -30$ , anterior $y \geq 0$ , mid otherwise). Up to three local maxima at least 8 mm apart, each exceeding all 26 neighboring voxels, are listed per cluster; $x$ , $y$ , $z$ are MNI coordinates. Sil, silence; Rev, reversed speech; Obj, object sentences; SE, social-emotional sentences; STG, superior temporal gyrus; STS, superior temporal sulcus; MTG, middle temporal gyrus; SMA, supplementary motor area.

**Figure 1.**
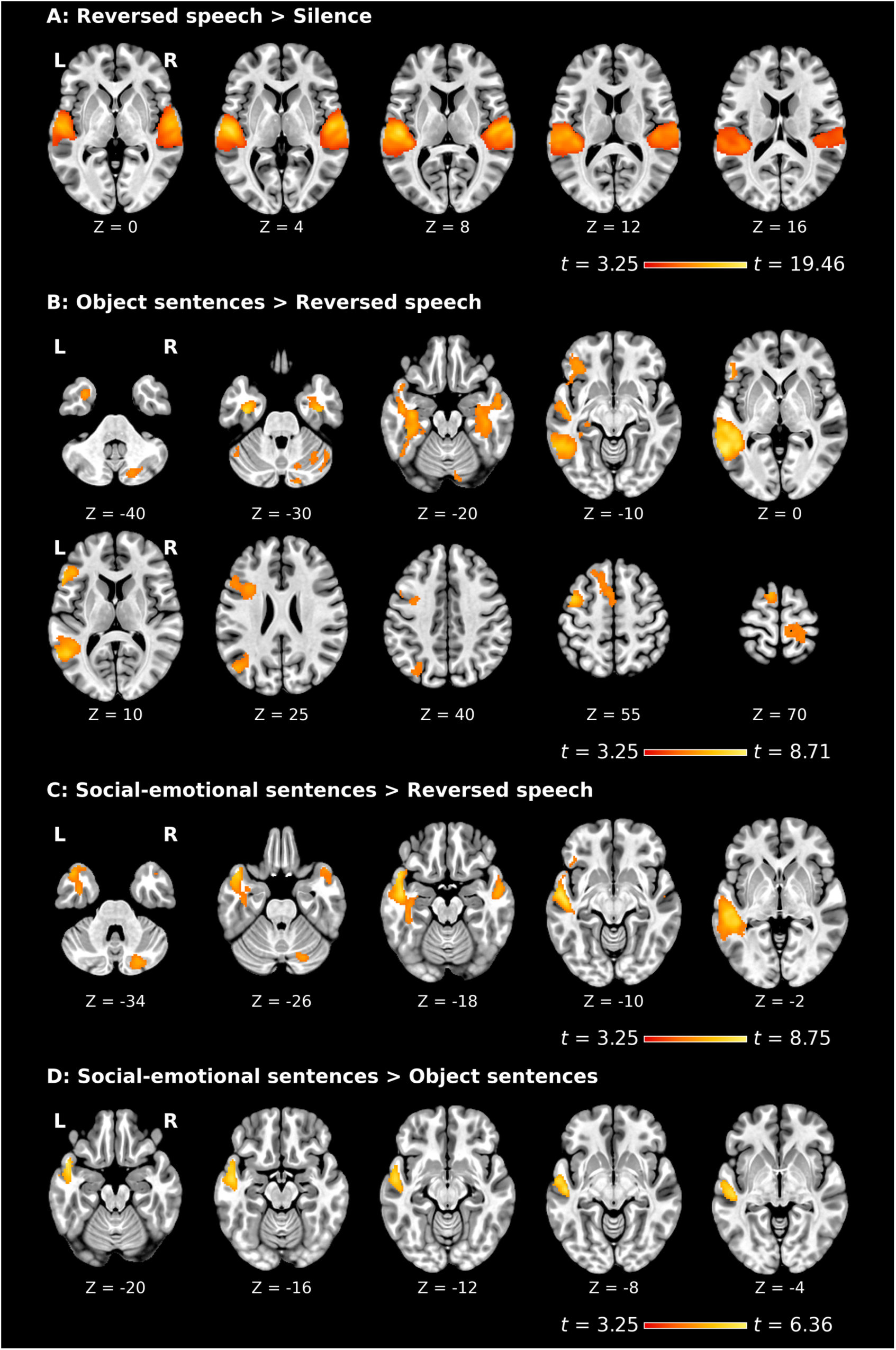
Whole-brain activity across all participants (*N* = 58). Second-level statistical maps for (A) Reversed speech > Silence, (B) Object sentences > Reversed speech, (C) Social-emotional sentences > Reversed speech, and (D) Social-emotional sentences > Object sentences. Clusters were formed at a voxel-level threshold of *p* < 0.001 (*t* = 3.25, *df* = 52) and are displayed only if they survived whole-brain cluster-level family-wise error correction at *p* < 0.05.

We also estimated every contrast within each diagnostic group separately. Both groups showed the expected pattern on the three contrasts that do not isolate social-emotional semantics (Figure S1, S2; Table S2a, S2b). Reversed speech relative to silence activated bilateral superior temporal cortex in each group; object sentences relative to reversed speech activated left temporal and left inferior frontal cortex in each group; and social-emotional sentences relative to reversed speech activated left superior temporal cortex in each group (Figure 2A). The groups diverged only on the contrast between the two sentence types (Figure 2B). In non-autistic children, [Social-emotional > Object] yielded a left STG/STS cluster (peak *t* = 8.94 at [−50, 2, −20], *k* = 641, *p* < 0.001), reproducing the pattern reported in non-autistic adults. In autistic children the same contrast yielded a cluster in the left superior temporal cortex (peak *t* = 4.55 at [−50, −18, −6], *k* = 36) that did not survive whole-brain correction (*p* = 0.833).

**Figure 2.**
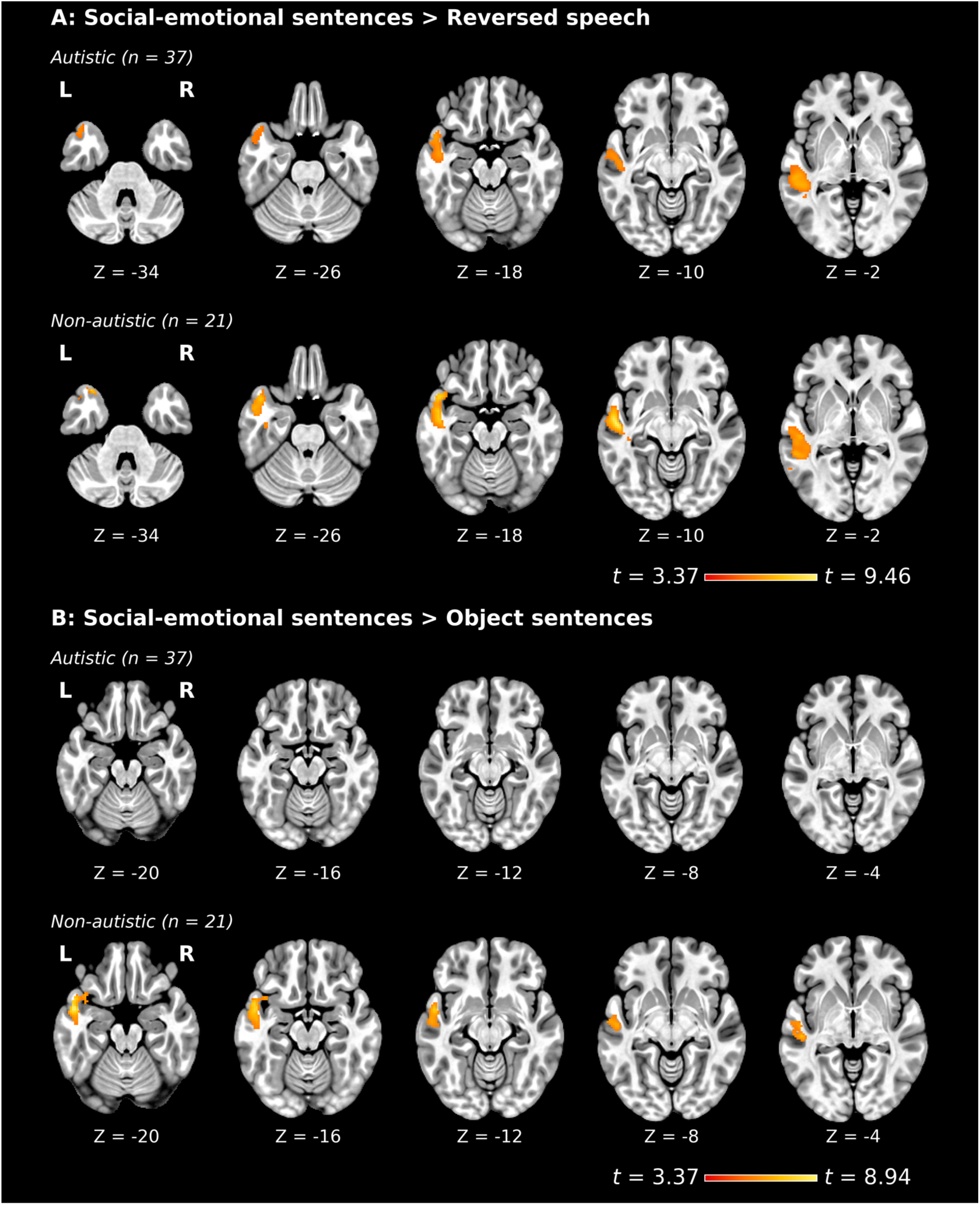
Social-emotional activation within autistic and non-autistic groups. (A) [Social-emotional sentences > Reversed speech] and (B) [Social-emotional sentences > Object sentences], estimated separately in the autistic (*n* = 37) and non-autistic (*n* = 21) groups with the same model and covariates as the across-group analysis. Clusters were formed at a voxel-level threshold of *p* < 0.001 (autistic *t* = 3.37, *df* = 32; non-autistic *t* = 3.69, *df* = 16) and are displayed only if they survived whole-brain cluster-level family-wise error correction at *p* < 0.05. In the autistic group no cluster survived correction for [Social-emotional > Object]; the largest sub-threshold cluster in that map (*k* = 36, peak *t* = 4.55, cluster-level *p* = 0.833) is listed in Table S2a.

We then compared the diagnostic groups at the whole brain, with group entered as a predictor. No contrast yielded a group difference surviving cluster-level FWE correction. This was true of the critical [Social-emotional > Object] contrast (largest cluster, non-autistic > autistic: *k* = 21, cluster-level *p* = 0.95; autistic > non-autistic: no cluster larger than one voxel) and of [Social-emotional > Reversed speech], and equally of the control contrasts, [Reversed speech > Silence] (largest *k* = 13, *p* = 0.99) and [Object sentences > Reversed speech] (largest *k* = 18, *p* = 0.97). This suggests that the groups did not differ in basic auditory or in general lexico-syntactic processing at the whole-brain level.

### ROI Analyses

At the a-priori left m-STS region taken from Mellem et al. (2016), autistic children showed significantly reduced activity compared to non-autistic children for the [Social-emotional > Object] contrast (autistic *M* = 0.13, non-autistic *M* = 0.48; *t* = 2.76, *p* = 0.008, FDR *p* = 0.031, *d* = 0.75). The same region showed a significant reduction for the [Social-emotional > Reversed speech] contrast as well (*t* = 2.85, *p* = 0.006, FDR *p* = 0.025, *d* = 0.78), whereas no other literature region reached significance in either contrast (Table 4, Figure 3B). Computing the same comparisons from unsmoothed data gave larger effects at the m-STS ([Social-emotional > Object]: *t* = 2.94, FDR *p* = 0.019; [Social-emotional > Reversed speech]: *t* = 3.20, FDR *p* = 0.009). At the left a-STG the difference for [Social-emotional > Reversed speech] was also larger without smoothing and reached significance there (*t* = 2.76, FDR *p* = 0.016) where it did not with it. At the a-STS and p-STS, where no difference was significant with smoothing, the effects were smaller without it and remained non-significant (Table S9). As a post-hoc test of the m-STS group difference, we examined whether each group individually showed a social-emotional bias, using one-sample *t* tests of the [Social-emotional > Object] contrast estimates against zero: the non-autistic group showed a significant bias (*t*(20) = 4.06, *p* < 0.001, *d* = 0.89), whereas the bias did not reach significance in the autistic group (*t*(36) = 1.91, *p* = 0.064, *d* = 0.31). The group difference was unchanged when parent-reported Cantonese or English proficiency, listening proficiency, or language dominance were added to the model (all *p* ≤ 0.012), and no language measure predicted m-STS activity (all *p* ≥ 0.65; Table S4).

**Table 4.**
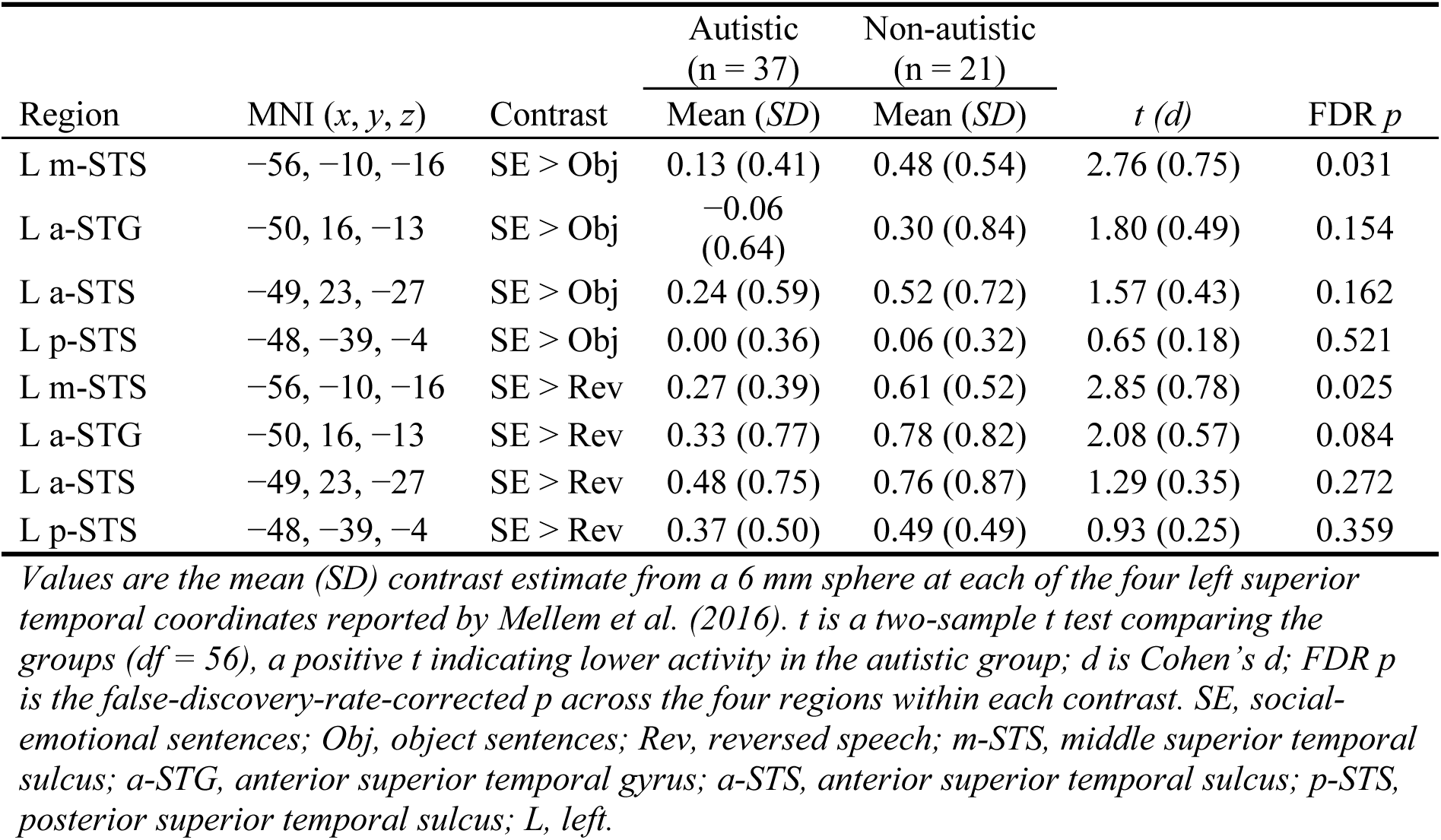
Group differences in region-of-interest activity between autistic and non-autistic children.

**Figure 3.**
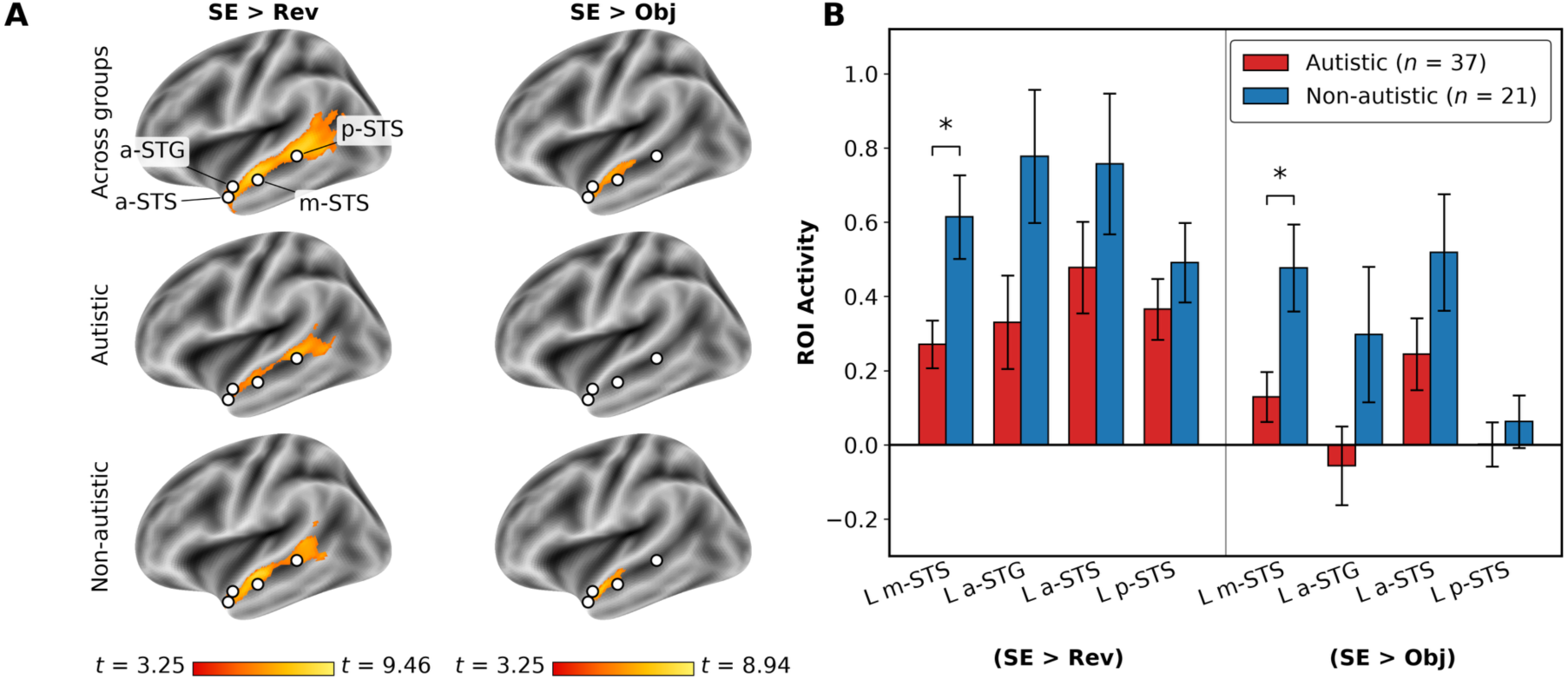
Left superior temporal response to social-emotional sentences, and activity at the a-priori regions of interest. (A) Statistical maps for [Social-emotional > Reversed speech] and [Social-emotional > Object sentences], estimated across all participants and separately within each diagnostic group, and shown on the left lateral surface only. Clusters were formed at a voxel-level threshold of *p* < 0.001 (across groups *t* = 3.25, *df* = 52; autistic *t* = 3.37, *df* = 32; non-autistic *t* = 3.69, *df* = 16) and are shown only if they survived whole-brain cluster-level family-wise error correction at *p* < 0.05. In the autistic group no cluster survived correction for [Social-emotional > Object]. White dots mark the four a-priori regions of interest of Mellem et al. (2016). (B) Mean contrast estimate for each group at every a-priori region of interest, extracted from a 6 mm sphere. Error bars are the standard error of the mean; an asterisk marks a group difference that is significant after false-discovery-rate correction (*p* < 0.05). SE, social-emotional sentences; Obj, object sentences; Rev, reversed speech; m-STS, middle superior temporal sulcus; a-STG, anterior superior temporal gyrus; a-STS, anterior superior temporal sulcus; p-STS, posterior superior temporal sulcus; L, left.

We next separated the social-emotional sentences by valence. Descriptively, the group difference was carried mainly by negative content, although, as set out below, the Group by Valence interaction was not significant. At the left m-STS, autistic children showed significantly reduced activity relative to non-autistic children for negative social-emotional sentences ([Social-emotional negative > Object]: autistic *M* = 0.06, non-autistic *M* = 0.49; *t* = 3.06, *p* = 0.003, FDR *p* = 0.013, *d* = 0.84), whereas for positive social-emotional sentences the difference ran in the same direction but was smaller and not significant (autistic *M* = 0.20, non-autistic *M* = 0.46; *t* = 1.75, *p* = 0.085, FDR *p* = 0.291, *d* = 0.48). The negative effect was also larger than the positive effect at the a-STG (*d* = 0.48 against *d* = 0.32), although it did not survive correction there (Table S5). A 2 (Valence: positive, negative) by 2 (Group: autistic, non-autistic) mixed ANOVA confirmed a significant main effect of Group at the m-STS (*F*(1,56) = 7.56, *p* = 0.008) but no significant Group by Valence interaction (*F*(1,56) = 1.20, *p* = 0.28).

We then explored whether the reduced m-STS bias varied continuously with autism symptom level or language ability. The group difference itself was robust to the full covariate set (β = −0.35, *p* = 0.010, *N* = 57) and remained significant when language ability and its interaction with group were entered (group simple effect at the mean HKCOLAS score: β = −0.45, *p* = 0.017, *N* = 50; interaction *p* = 0.45; HKCOLAS was available for 29 of 37 autistic children and all 21 non-autistic children), indicating that it was not explained by language ability. It could not, however, be statistically separated from autism symptom level: diagnostic group and ADOS-2 score are collinear by construction (*r* = 0.88) and their ranges do not overlap, so that entering the ADOS-2 total and its interaction with group left neither the group simple effect (*p* = 0.56) nor the ADOS-2 term (*p* = 0.86) significant, and the data are equally compatible with a diagnostic and with a continuous symptom-level account of the difference. Within each group, however, we found no evidence of a continuous relationship: neither ADOS-2 total scores nor HKCOLAS predicted m-STS activity in the autistic group (both p ≥ 0.74) or in the non-autistic group (both *p* ≥ 0.23), no predictor-by-group interaction was significant, and the within-group tests for the ADOS-2 social-affect and restricted-repetitive-behavior subscales were likewise non-significant (Table S6, Figure S3). These within-group analyses are underpowered for the small effects that continuous brain-behavior associations typically involve, so we treat the absence of a gradient as suggestive. Age did not predict left m-STS activity across the sample (*r* = 0.04, *p* = 0.79) or within either group (both *p* > 0.46).

## DISCUSSION

The current study investigated the neural mechanisms underlying the processing of social-emotional meaning in language among Cantonese-speaking autistic and non-autistic children. We found that the left STG/STS showed a neural bias for social-emotional sentences over object sentences and reversed speech. This bias is in line with what Mellem et al. (2016) reported in non-autistic adults, and it shows that the social-emotional bias of the left STG/STS is already present in children aged 8 to 12.5. Consistent with our hypothesis, this bias was significantly reduced in autistic children compared with non-autistic peers. This reduced neural bias we observe in autistic children is, to our knowledge, the first demonstration that the social-emotional bias of the left STG/STS is altered in autism at the level of semantics, beyond the level of paralinguistic or non-verbal social cues.

The group difference in the left m-STS was not accompanied by detectable differences in general auditory or lexico-syntactic processing: no group difference survived correction in the [Reversed speech > Silence] contrast or in the [Object sentences > Reversed speech] contrast. Individual-difference analyses indicated that the reduced bias is not explained by language ability: the group difference persisted after adjusting for HKCOLAS scores, and language ability did not predict STG/STS activity within either group. It could not, however, be statistically separated from autism symptom level, because diagnostic group and ADOS-2 scores are collinear by construction. What the data do show is that no gradient of symptom level was detectable within the autistic group itself.

A further implication concerns the level of language processing at which these differences emerge. Communicative challenges in autism have been reported most often at the pragmatic level, that is, the use and interpretation of language in social context (Loukusa & Moilanen, 2009; Schaeffer et al., 2023). Difficulties at the level of semantic processing, by contrast, have been documented less consistently: reviews of lexical semantic ability in autism report roughly equal numbers of studies finding comparable and lower performance, with no single factor explaining the variability (Sukenik & Tuller, 2023), and language profiles are highly heterogeneous across autistic individuals more broadly (Boucher, 2012; Kjelgaard & Tager-Flusberg, 2001). Our results speak to this inconsistency by pointing to a more specific locus. Object sentences, which convey non-social meaning, elicited comparable responses in the two groups, whereas the response to sentences conveying social and emotional meaning was reduced in autistic children at the left m-STS. Within the autistic group, responses to reversed speech and to object sentences were robust, while [Social-emotional > Object] produced no cluster surviving correction (Figure 2, Table S2a). What distinguishes the two groups may therefore not be semantic processing in general, but the social-emotional subset of it, which extends the language differences documented in autism from the pragmatic level down to the semantics of social and emotional meaning.

The reduced activity we observe in the left STG/STS offers a potential neural mechanism for a pattern described in the autism literature. Autistic adults can access the social knowledge that speech conveys but appear to do so less automatically and with greater effort (Tesink et al., 2009). Evidence from tasks requiring explicit judgments is nonetheless mixed, with roughly half of them reporting group differences (Lartseva et al., 2015). The left STG/STS has been suggested to be a multimodal interface that parses dynamic input across modalities and recovers the communicative meaning that input carries (Kausel et al., 2024; Redcay, 2008). It is further suggested that in autism the dysfunction of this region may affect the automatic recovery of meaning more than the parsing itself, although deficits in processing rapidly changing input have also been reported. Our results align with this account and extend its scope. Previous work has grounded it in non-linguistic visual cues, such as biological motion and eye gaze, and in paralinguistic auditory cues, such as prosody and vocal intonation; we show the same signature for meaning carried by the language content itself. The reduced recruitment of the left STG/STS therefore appears to be organized around the social and emotional meaning rather than the channel that conveys it, whether a gesture, a tone of voice, or the words themselves.

The reduced social-emotional bias in autism was carried mainly by sentences with negative emotional content, for which the group difference was significant (*d* = 0.84), whereas positive sentences showed a smaller, non-significant difference in the same direction (*d* = 0.48). Because the Group by Valence interaction was not significant, we treat this pattern as suggestive. It nonetheless converges with reports that autistic traits are associated with altered processing of social-negative emotional content (Yang et al., 2022), and it indicates that the effect we observe might not be driven uniformly by all emotional content.

Several limitations should be noted. First, the final sample included for analysis was modest by current neuroimaging standards, particularly the non-autistic group. A sensitivity analysis, computed with the pwr package (version 1.3-0; Champely, 2020) in R version 4.3.2 (R Core Team, 2023), with two-tailed α = 0.05 and 37 versus 21 participants, indicated that the sample had 80% power only for between-group effects of Cohen’s *d* = 0.78 or larger. The reductions we observed at the left m-STS (*d* = 0.75 and *d* = 0.78) sit at the edge of what this design was well placed to detect. Smaller effects, including the positive-valence contrast (*d* = 0.48), were beyond its resolution, so those null results are inconclusive rather than evidence of no difference. This limitation applies with particular force to the individual-difference analyses, since continuous brain-behavior associations tend to be small and their reliable detection generally requires samples far larger than ours (Marek et al., 2022). The absence of a within-group gradient in symptom level or language ability should therefore be read as suggestive rather than conclusive. Second, although the sex distribution did not differ between groups and sex was included as a predictor in all whole-brain models, the sample was predominantly male (48 of 58 participants), reflecting the higher rate of autism diagnosis in boys. Therefore, the present findings may not generalize equally to autistic girls. Sex-balanced samples will be needed to establish whether the reduced social-emotional bias observed here is comparable across sexes. Third, head motion excluded a larger proportion of the autistic children than of the non-autistic children (31% against 13%), so the autistic children in the final sample are those able to lie still throughout the scan, and the findings may not extend to autistic children with greater head motion. Within the analyzed sample, residual motion was closely matched (mean framewise displacement 0.25 vs. 0.25, *p* = 0.92; high-motion volumes 6.65% vs. 7.08%, *p* = 0.80). A further limitation concerns the social-emotional contrast: because the object sentences were non-social and affectively neutral, it varies social content together with emotional valence and arousal and cannot fully separate them. Separating them would require the social but affectively neutral condition.

In conclusion, the left STG/STS is more responsive to language that carries social and emotional meaning than to language that does not, in children as in adults, and this neural bias is reduced in autistic children. This finding suggests that the social communication difficulties in autism may stem not only from challenges in reading non-verbal or paralinguistic cues but also from altered processing of social and emotional content at the semantic level. However, it should be acknowledged that the current study, serving as a starting point, focused exclusively on regional activity patterns. The reduced activity in the left STG/STS in autism could be a downstream consequence of network connectivity differences. Future research should investigate whether the reduced activity of the left STG/STS stems from its disconnection from the broader social brain network.

## ACKNOWLEDGMENTS

This work was supported by Health and Medical Research Fund, Hong Kong (HMRF 10211176) and the National Social Science Fund of China (22CYY021). We thank the three anonymous reviewers for their constructive comments, and the audience at the International Symposium on Language Science (ISLS) 2026, where this work was presented, for their questions and helpful discussion.

