## Supplementary materials for "Autistic children show reduced activity in left superior temporal cortex in response to social-emotional meaning in language"

**Table S1.** Experimental materials with English translation

| Type | Sentence | English translation | MWF | MDD | WC |
|---|---|---|---|---|---|
| Social-emotional | | | | | |
| Positive | 醫生收到好多誇獎。 | The doctor received many compliments. | 3.34 | 2.33 | 6 |
| | 老師經常讚同學叻。 | The teacher often praises the students for being smart. | 1.57 | 2.40 | 5 |
| | 班長對人好有禮貌。 | The class monitor is very polite to everyone. | 2.80 | 2.83 | 6 |
| | 船長對自己有信心。 | The captain has confidence in himself. | 2.21 | 2.60 | 5 |
| | 社工幫咗好多學生。 | The social worker helped a lot of students. | 3.34 | 2.33 | 6 |
| | 公主係個快樂嘅人。 | The princess is a happy person. | 3.20 | 3.00 | 6 |
| | 公公見到彩虹好靚。 | Grandpa saw a beautiful rainbow. | 2.86 | 2.50 | 6 |
| | 哥哥好錫佢女朋友。 | My older brother loves his girlfriend very much. | 3.00 | 2.40 | 5 |
| | 婆婆行街行到過癮。 | Grandma thoroughly enjoyed shopping. | 2.28 | 2.20 | 5 |
| | 姐姐聽笑話聽到笑。 | My older sister laughed at the joke. | 2.88 | 2.33 | 6 |
| | 小明有好好嘅朋友。 | Siu-Ming has very good friends. | 3.63 | 2.67 | 6 |
| | 小明有好大嘅本事。 | Siu-Ming is very capable. | 3.56 | 2.67 | 6 |
| | 小明好有英雄氣概。 | Siu-Ming has a heroic spirit. | 2.77 | 2.40 | 5 |
| | 嘉琪成功咗好激動。 | Ka-Kei was thrilled by her success. | 2.91 | 2.60 | 5 |
| | 俊俊恭喜結婚嘅人。 | Jun-Jun congratulated the newlyweds. | 2.70 | 2.60 | 5 |
| | 俊俊興奮到跳起身。 | Jun-Jun was so excited that he jumped up. | 2.21 | 3.17 | 6 |
| | 琳琳好熱心咁幫人。 | Lam-Lam is very eager to help others. | 3.13 | 2.50 | 6 |
| | 超人覺得自己好勁。 | Superman thinks he is very powerful. | 2.62 | 2.60 | 5 |
| Negative | 司機太心急撞咗車。 | The driver crashed the car in haste. | 2.36 | 2.67 | 6 |
| | 魔鬼個樣好得人驚。 | The devil looked terrifying. | 2.43 | 2.20 | 5 |
| | 爺爺跌咗錢好心痛。 | Grandpa was heartbroken over losing money. | 3.24 | 2.67 | 6 |
| | 爸爸屙嘅大便好臭。 | Dad's poop was very smelly. | 3.23 | 2.17 | 6 |
| | 爸爸後悔冇早啲走。 | Dad regretted not leaving earlier. | 2.78 | 2.33 | 6 |
| | 爺爺鄙視粗暴嘅人。 | Grandpa despises violent people. | 1.76 | 2.60 | 5 |

**Table S1.** Experimental materials with English translation

| Type | Sentence | English translation | MWF | MDD | WC |
|---|---|---|---|---|---|
| Social-emotional | | | | | |
| | 細佬考試考到緊張。 | My little brother got nervous taking the exam. | 1.69 | 2.20 | 5 |
| | 媽媽有好多嘅煩惱。 | Mom has so many worries. | 3.69 | 3.00 | 6 |
| | 志強覺得自己好差。 | Chi-Keung feels he is no good. | 2.00 | 2.60 | 5 |
| | 志強諗起啲傷心事。 | Chi-Keung recalled some sad memories. | 2.15 | 2.50 | 6 |
| | 玲玲覺得課室好嘈。 | Ling-Ling thinks the classroom is very noisy. | 2.04 | 2.20 | 5 |
| | 家樂等到好唔耐煩。 | Ka-Lok grew very impatient while waiting. | 3.10 | 3.00 | 6 |
| | 媽媽覺得爸爸好煩。 | Mom finds dad very annoying. | 2.99 | 2.60 | 5 |
| | 媽媽好惡咁鬧細佬。 | Mom scolded my little brother very fiercely. | 2.93 | 3.00 | 6 |
| | 受傷嘅人望落好慘。 | The injured person looked miserable. | 2.77 | 2.33 | 6 |
| | 細佬見到蜘蛛就驚。 | My little brother gets scared whenever he sees a spider. | 3.08 | 2.83 | 6 |
| | 媽媽覺得細妹好惡。 | Mom thinks my little sister is very mean. | 2.39 | 2.60 | 5 |
| | 家樂覺得細佬好蠢。 | Ka-Lok thinks his little brother is very stupid. | 2.40 | 2.60 | 5 |
| Object | | | | | |
| | 正方盒入面有粒波。 | There is a ball inside the square box. | 2.96 | 2.20 | 5 |
| | 生果盆入面有粒橙。 | There is an orange in the fruit bowl. | 2.92 | 2.20 | 5 |
| | 雨水由屋頂流落地。 | Rainwater flows from the roof to the ground. | 2.34 | 2.67 | 6 |
| | 地下嘅磚係紅色嘅。 | The floor tiles are red. | 3.02 | 2.17 | 6 |
| | 牆上面黐咗張貼紙。 | There is a sticker stuck on the wall. | 2.33 | 2.33 | 6 |
| | 鞋帶綁咗喺鞋上面。 | The shoelaces are tied on the shoes. | 2.71 | 2.67 | 6 |
| | 窗上面有一塊窗簾。 | There is a curtain on the window. | 2.46 | 2.33 | 6 |
| | 地球圍住太陽轉圈。 | The Earth revolves around the Sun. | 1.89 | 2.20 | 5 |
| | 菠蘿嘅肉係黃色嘅。 | The flesh of a pineapple is yellow. | 3.02 | 2.17 | 6 |
| | 一年會有四個季節。 | There are four seasons in a year. | 3.39 | 2.43 | 7 |
| | 光可以穿過玻璃樽。 | Light can pass through a glass bottle. | 2.76 | 2.40 | 5 |

**Table S1.** Experimental materials with English translation

| Type | Sentence | English translation | MWF | MDD | WC |
|---|---|---|---|---|---|
| Social-emotional | | | | | |
| | 紙係由木頭做成嘅。 | Paper is made from wood. | 2.70 | 2.57 | 7 |
| | 檯上面有兩本雜誌。 | There are two magazines on the table. | 2.42 | 2.50 | 6 |
| | 蘋果熟咗會跌落地。 | Apples fall to the ground when they are ripe. | 2.99 | 2.43 | 7 |
| | 月亮本身唔會發光。 | The moon does not produce its own light. | 2.46 | 3.20 | 5 |
| | 橋嘅下面有一條河。 | There is a river under the bridge. | 2.88 | 2.43 | 7 |
| | 太陽係由東邊升起。 | The sun rises in the east. | 2.39 | 2.86 | 7 |
| | 山上面有個大山窿。 | There is a large cave on the mountain. | 2.87 | 2.50 | 6 |
| | 麵包係用麵粉做嘅。 | Bread is made from flour. | 2.94 | 2.50 | 6 |
| | 湯同茶係一種液體。 | Soup and tea are liquids. | 2.97 | 2.43 | 7 |
| | 路邊有幾舊小石頭。 | There are a few small stones by the roadside. | 2.74 | 2.67 | 6 |
| | 粥係用米同水煮成。 | Congee is made by boiling rice and water. | 2.78 | 3.25 | 8 |
| | 樹葉係樹嘅一部分。 | Leaves are part of a tree. | 3.11 | 2.83 | 6 |
| | 泥土濕咗會變深色。 | Soil becomes darker when it is wet. | 2.52 | 2.33 | 6 |
| | 樹根埋喺泥土下面。 | Tree roots lie buried beneath the soil. | 2.33 | 2.20 | 5 |
| | 一日有廿四個小時。 | There are twenty-four hours in a day. | 2.98 | 2.33 | 6 |
| | 水燒到一百度會滾。 | Water boils when heated to 100 degrees. | 2.72 | 3.14 | 7 |
| | 光照到鏡就會反射。 | Light reflects when it hits a mirror. | 2.74 | 3.29 | 7 |
| | 水燒滾咗會有泡泡。 | Boiling water has bubbles. | 2.72 | 2.29 | 7 |
| | 英文有廿六個字母。 | There are twenty-six letters in the English alphabet. | 2.94 | 2.40 | 5 |
| | 三角形係一個圖案。 | A triangle is a shape. | 2.84 | 2.40 | 5 |
| | 樹葉生喺樹枝上面。 | Leaves grow on branches. | 1.78 | 2.20 | 5 |
| | 海水曬乾會變成鹽。 | Seawater turns into salt when dried. | 2.09 | 2.29 | 7 |
| | 香蕉嘅皮係黃色嘅。 | Banana peels are yellow. | 3.15 | 2.17 | 6 |
| | 秋天嘅樹葉會變黃。 | Leaves turn yellow in autumn. | 2.62 | 2.33 | 6 |
| | 荔枝嘅殼係紅色嘅。 | Lychee shells are red. | 2.68 | 2.17 | 6 |

*MWF, mean word frequency; MDD, mean dependency distance; WC, word count.*

**Table S2a.** Clusters and local maxima with significant activity in the autistic group (n = 37)

| Contrast | Region | Cluster size | *x* | *y* | *z* | *t* | *p* |
|---|---|---|---|---|---|---|---|
| Rev > Sil | L STG (auditory cortex) | 2205 | −54 | −22 | 8 | 16.77 | < 0.001 |
| | | | −40 | −32 | 10 | 10.09 | |
| | | | −66 | −24 | 10 | 9.87 | |
| | R STG (auditory cortex) | 2054 | 56 | −14 | 4 | 13.52 | < 0.001 |
| | | | 68 | −20 | 6 | 10.77 | |
| | | | 60 | −2 | 0 | 9.92 | |
| Obj > Rev | L STS (posterior) | 3610 | −56 | −50 | 0 | 7.12 | < 0.001 |
| | | | −48 | −48 | −2 | 6.68 | |
| | | | −50 | −40 | 2 | 6.53 | |
| | R fusiform gyrus | 875 | 38 | −8 | −32 | 5.87 | < 0.001 |
| | | | 32 | −12 | −18 | 5.21 | |
| | | | 48 | −6 | −20 | 5.07 | |
| | L precentral gyrus | 405 | −42 | 4 | 52 | 5.95 | < 0.001 |
| | | | −36 | −8 | 60 | 4.46 | |
| | | | −32 | −4 | 66 | 3.61 | |
| | R postcentral gyrus | 384 | 14 | −34 | 78 | 4.41 | < 0.001 |
| | | | 26 | −36 | 74 | 4.21 | |
| | | | 8 | −32 | 70 | 4.1 | |
| | L IFG (pars triangularis) | 366 | −56 | 20 | 12 | 5.58 | 0.001 |
| | | | −54 | 28 | 8 | 4.78 | |
| | | | −46 | 30 | 12 | 4.6 | |
| | L SMA | 198 | −8 | 12 | 64 | 5.23 | 0.028 |
| | | | −10 | 4 | 68 | 4.65 | |
| | | | −10 | 6 | 52 | 4.14 | |
| SE > Rev | L STS (posterior) | 1748 | −54 | −34 | 0 | 6.69 | < 0.001 |
| | | | −58 | −56 | 6 | 5.75 | |
| | | | −52 | 10 | −20 | 5.51 | |
| SE > Obj | L STS (mid) | 36 | −50 | −18 | −6 | 4.55 | 0.833 (n.s.) |

*The autistic group was analyzed on its own, with the same model and covariates as the across-group analysis (sex, age, mean framewise displacement, non-verbal IQ). Clusters were formed at a voxel-level threshold of* $p < 0.001$ *(*$t = 3.37$*,* $df = 32$*) and retained only if they survived whole-brain cluster-level family-wise error correction at* $p < 0.05$*; p is the cluster-level FWE-corrected value. Labeling, local-maxima selection and coordinates follow Table 3 of the main text. For [Social-emotional > Object] no cluster survived correction; the largest sub-threshold cluster is listed for completeness and marked (n.s.). Sil, silence; Rev, reversed speech; Obj, object sentences; SE, social-emotional sentences; STG, superior temporal gyrus; STS, superior temporal sulcus; SMA, supplementary motor area; IFG, inferior frontal gyrus.*

**Table S2b.** Clusters and local maxima with significant activity in the non-autistic group (n = 21)

| Contrast | Region | Cluster size | *x* | *y* | *z* | *t* | *p* |
|---|---|---|---|---|---|---|---|
| Rev > Sil | L STG (auditory cortex) | 1701 | −60 | −14 | 2 | 12.31 | < 0.001 |
| | | | −42 | −24 | 12 | 9.15 | |
| | | | −38 | −34 | 8 | 8.21 | |
| | R STG (auditory cortex) | 1505 | 54 | −14 | 6 | 11.18 | < 0.001 |
| | | | 66 | −10 | 4 | 8.7 | |
| | | | 56 | −6 | 2 | 8.37 | |
| Obj > Rev | L STS (posterior) | 1290 | −56 | −32 | 0 | 7.86 | < 0.001 |
| | | | −58 | −56 | 0 | 6.92 | |
| | | | −42 | −46 | 2 | 6.87 | |
| | R cerebellum (lobule VIII) | 169 | 28 | −66 | −50 | 5.55 | 0.020 |
| | | | 20 | −76 | −44 | 5.54 | |
| | L IFG (pars triangularis) | 165 | −50 | 36 | 0 | 4.96 | 0.023 |
| | | | −50 | 24 | 4 | 4.95 | |
| | | | −48 | 18 | −6 | 4.2 | |
| SE > Rev | L STS (mid) | 2031 | −54 | −12 | −10 | 9.46 | < 0.001 |
| | | | −50 | −2 | −20 | 7.36 | |
| | | | −48 | 14 | −18 | 7.26 | |
| SE > Obj | L STS (anterior) | 641 | −50 | 2 | −20 | 8.94 | < 0.001 |
| | | | −52 | 10 | −18 | 7.13 | |
| | | | −54 | 2 | −12 | 6.73 | |

*The non-autistic group was analyzed on its own, with the same model and covariates as the across-group analysis (sex, age, mean framewise displacement, non-verbal IQ). Clusters were formed at a voxel-level threshold of* $p < 0.001$ *(*$t = 3.69$, $df = 16$*) and retained only if they survived whole-brain cluster-level family-wise error correction at* $p < 0.05$*; p is the cluster-level FWE-corrected value. Labeling, local-maxima selection and coordinates follow Table 3 of the main text. Sil, silence; Rev, reversed speech; Obj, object sentences; SE, social-emotional sentences; STG, superior temporal gyrus; STS, superior temporal sulcus; IFG, inferior frontal gyrus.*

**Table S3.** Language proficiency of autistic and non-autistic children

| Variable | | Autistic<br>Mean (SD) | Non-autistic<br>Mean (SD) | *t* | *df* | *p* |
|---|---|---|---|---|---|---|
| English | listening | 4.14 (1.57) | 4.57 (1.47) | -1.06 | 43.97 | 0.294 |
| | speaking | 3.89 (1.54) | 4.67 (1.15) | -2.17 | 51.60 | 0.035 |
| | reading | 3.97 (1.61) | 4.90 (1.09) | -2.62 | 54.05 | 0.011 |
| | writing | 3.78 (1.55) | 4.38 (1.16) | -1.66 | 51.56 | 0.102 |
| | composite | 3.95 (1.48) | 4.63 (1.06) | -2.04 | 52.81 | 0.047 |
| Cantonese | listening | 6.38 (1.04) | 6.76 (0.44) | -1.96 | 52.73 | 0.055 |
| | speaking | 6.22 (1.03) | 6.71 (0.46) | -2.52 | 53.88 | 0.015 |
| | reading | 5.19 (1.54) | 5.86 (1.15) | -1.87 | 51.67 | 0.067 |
| | writing | 4.81 (1.68) | 5.43 (1.36) | -1.52 | 49.10 | 0.134 |
| | composite | 5.65 (1.06) | 6.19 (0.71) | -2.32 | 54.28 | 0.024 |
| Dominance | composite | 1.70 (1.52) | 1.56 (1.29) | 0.38 | 47.47 | 0.705 |
| | listening | 2.24 (1.85) | 2.19 (1.57) | 0.12 | 47.51 | 0.909 |

*Proficiency was rated by parents on a scale from 0 to 7 for each dimension; reading and writing refer to written Chinese, the literacy form used by Cantonese speakers. Composite = mean of the listening, speaking, reading, and writing ratings; Dominance = Cantonese minus English proficiency. Group comparisons are Welch (unequal-variance) t tests (37 autistic, 21 non-autistic); a positive t indicates a higher rating in the autistic group.*

**Table S4.** Sensitivity of the left m-STS group difference to language experience

| Language measure added to the model | Group effect | | | Language measure | | *N* |
|---|---|---|---|---|---|---|
| | β | *t* | *p* | β | *p* | |
| (none: base model) | -0.347 | -2.68 | 0.010 | | | 57 |
| Cantonese composite proficiency | -0.367 | -2.66 | 0.010 | -0.030 | 0.654 | 57 |
| English composite proficiency | -0.354 | -2.61 | 0.012 | -0.008 | 0.855 | 57 |
| Cantonese listening | -0.362 | -2.63 | 0.011 | -0.026 | 0.727 | 57 |
| English listening | -0.354 | -2.68 | 0.010 | -0.014 | 0.732 | 57 |
| Cantonese-English dominance (composite) | -0.347 | -2.65 | 0.011 | -0.005 | 0.908 | 57 |
| Cantonese-English dominance (listening) | -0.347 | -2.65 | 0.011 | +0.005 | 0.892 | 57 |

*Each row is a linear regression of the left m-STS [Social-emotional > Object] contrast estimate on diagnostic group with the standard covariate set (age, sex, mean framewise displacement, non-verbal IQ), plus one parent-reported language measure; the first row is the base model with no language measure. Group β is coded autistic minus non-autistic, so a negative value indicates lower activity in autistic children. All continuous predictors were mean-centered.*

**Table S5.** Group differences in social-emotional activity separated by valence

| Region | Contrast | Autistic | Non-autistic | *t* | *d* | FDR *p* |
|---|---|---|---|---|---|---|
| L m-STS (−56, −10, −16) | SE positive > Object | 0.20 (0.59) | 0.46 (0.50) | 1.75 | 0.48 | 0.291 |
| | SE negative > Object | 0.06 (0.41) | 0.49 (0.65) | 3.06 | 0.84 | 0.013 |
| L a-STG (−50, 16, −13) | SE positive > Object | −0.09 (0.90) | 0.17 (0.65) | 1.16 | 0.32 | 0.334 |
| | SE negative > Object | −0.04 (0.81) | 0.42 (1.19) | 1.76 | 0.48 | 0.169 |
| L a-STS (−49, 23, −27) | SE positive > Object | 0.16 (0.85) | 0.49 (0.82) | 1.48 | 0.40 | 0.291 |
| | SE negative > Object | 0.33 (0.76) | 0.56 (0.97) | 0.98 | 0.27 | 0.333 |
| L p-STS (−48, −39, −4) | SE positive > Object | −0.03 (0.54) | −0.05 (0.33) | −0.11 | −0.03 | 0.912 |
| | SE negative > Object | 0.04 (0.38) | 0.18 (0.49) | 1.20 | 0.33 | 0.313 |

*Contrast estimates were extracted from 6 mm spheres at the four left superior temporal coordinates of Mellem et al. (2016). t is a two-sample test comparing the groups (df = 56; positive t indicates lower activity in the autistic group); FDR p is corrected across the four regions within each valence contrast. SE, social-emotional sentences.*

**Table S6.** Linear regression models of left m-STS [Social-emotional > Object] activity

| Model | Predictor | *β* | *t* | *p* | *N* |
|---|---|---|---|---|---|
| **Group difference** | | | | | |
| ROI ~ Group | Group (autistic − non-autistic) | -0.347 | -2.68 | .010 | 57 |
| **Autistic-trait level (ADOS-2 total)** | | | | | |
| Within autistic group | ADOS-2 total | +0.007 | 0.33 | .745 | 36 |
| Within non-autistic group | ADOS-2 total | -0.004 | -0.04 | .968 | 21 |
| Pooled (whole sample) | ADOS-2 total | -0.024 | -2.10 | .040 | 57 |
| Combined | ADOS-2 total | -0.013 | -0.18 | .860 | 57 |
| Combined | Group (autistic − non-autistic) | -0.295 | -0.59 | .559 | 57 |
| Combined | ADOS-2 total × Group | +0.022 | 0.29 | .770 | 57 |
| **Language ability (HKCOLAS)** | | | | | |
| Within autistic group | HKCOLAS | -0.002 | -0.14 | .893 | 29 |
| Within non-autistic group | HKCOLAS | -0.057 | -1.24 | .233 | 21 |
| Pooled (whole sample) | HKCOLAS | +0.008 | 0.57 | .570 | 50 |
| Combined | HKCOLAS | -0.029 | -0.96 | .343 | 50 |
| Combined | Group (autistic − non-autistic) | -0.454 | -2.50 | .017 | 50 |
| Combined | HKCOLAS × Group | +0.025 | 0.75 | .455 | 50 |
| **ADOS-2 social affect (secondary)** | | | | | |
| Within autistic group | ADOS-2 SA | +0.010 | 0.45 | .653 | 36 |
| Within non-autistic group | ADOS-2 SA | -0.088 | -0.96 | .352 | 21 |
| Pooled (whole sample) | ADOS-2 SA | -0.027 | -2.11 | .040 | 57 |
| Combined | ADOS-2 SA | -0.081 | -1.14 | .259 | 57 |
| Combined | Group (autistic − non-autistic) | +0.076 | 0.18 | .858 | 57 |
| Combined | ADOS-2 SA × Group | +0.092 | 1.23 | .224 | 57 |
| **ADOS-2 restricted/repetitive behavior (secondary)** | | | | | |
| Within autistic group | ADOS-2 RRB | -0.018 | -0.33 | .744 | 36 |
| Within non-autistic group | ADOS-2 RRB | +0.269 | 1.75 | .100 | 21 |
| Pooled (whole sample) | ADOS-2 RRB | -0.042 | -0.93 | .356 | 57 |
| Combined | ADOS-2 RRB | +0.251 | 1.90 | .064 | 57 |
| Combined | Group (autistic − non-autistic) | -0.637 | -3.15 | .003 | 57 |
| Combined | ADOS-2 RRB × Group | -0.263 | -1.82 | .076 | 57 |

*All models include age, sex, mean framewise displacement, and non-verbal IQ as covariates (continuous predictors mean-centered). ROI = 6 mm sphere at the Mellem m-STS coordinate (−56, −10, −16). Group is coded autistic minus non-autistic. N varies with missing data within the 58-child sample: eight autistic children lacked the HKCOLAS, and one of those eight also lacked the TONI-4.*

**Table S7.** Co-occurring diagnoses in the final sample

| Co-occurring diagnosis | Autistic (n = 37) | Non-autistic (n = 21) |
|---|---|---|
| Attention-deficit/hyperactivity disorder | 22 (59%) | 0 (0%) |
| Dyslexia | 8 (22%) | 0 (0%) |
| Developmental language or communication disorder | 8 (22%) | 0 (0%) |
| Anxiety disorder | 4 (11%) | 0 (0%) |
| Gastrointestinal disorder | 3 (8%) | 0 (0%) |
| Sleep disorder | 2 (5%) | 0 (0%) |
| Depression | 1 (3%) | 0 (0%) |
| Epilepsy | 1 (3%) | 0 (0%) |
| **Any co-occurring diagnosis** | **27 (73%)** | **0 (0%)** |
| **No co-occurring diagnosis** | **10 (27%)** | **21 (100%)** |

*Values are the number (percentage) of children with each diagnosis.*

**Table S8.** Sensitivity analysis: results with the participant lacking non-verbal IQ excluded

| Analysis | Measure | Main analysis (N = 58) | Sensitivity (N = 57) |
|---|---|---|---|
| Whole-brain [Social-emotional > Object] main effect | Peak (MNI) | −52, −18, −6 | −52, −18, −6 |
| | Peak *t* | 6.36 | 6.21 |
| | Cluster size (k) | 993 | 1013 |
| | Cluster-level *FWE p* | < 0.001 | < 0.001 |
| Whole-brain group comparison [Social-emotional > Object] | Autistic > non-autistic | No cluster survived | No cluster survived |
| | Non-autistic > autistic | No cluster survived | No cluster survived |
| Within-group whole-brain [Social-emotional > Object] | Non-autistic: k, peak *t*, *FWE p* | 641, 8.94, < 0.001 | 641, 8.94, < 0.001 |
| | Autistic: k, peak t, *FWE p* | 36, 4.55, 0.833 (n.s.) | 30, 4.33, 0.892 (n.s.) |
| Left m-STS ROI, group difference [Social-emotional > Object] | *t* (*df* = 56 / 55) | 2.76 | 2.67 |
| | *p* | 0.008 | 0.010 |

*The main analyses retain all 58 children and mean-impute the single missing TONI-4 value (one autistic child); the sensitivity analysis excludes that child (complete case, N = 57: 36 autistic, 21 non-autistic). Whole-brain values are the largest cluster (k, peak t, cluster-level FWE p; voxel-forming p < 0.001).*

**Table S9.** Group differences at the literature regions computed from unsmoothed data

| Contrast | Region | Smoothed: *t*, *FDR p* | Unsmoothed: *t* (*d*), *FDR p* |
|---|---|---|---|
| SE > Obj | L m-STS | 2.76, 0.031 | 2.94 (0.80), 0.019 |
| | L a-STG | 1.80, 0.154 | 2.04 (0.56), 0.092 |
| | L a-STS | 1.57, 0.162 | 0.93 (0.25), 0.474 |
| | L p-STS | 0.65, 0.521 | 0.28 (0.08), 0.780 |
| SE > Rev | L m-STS | 2.85, 0.025 | 3.20 (0.87), 0.009 |
| | L a-STG | 2.08, 0.084 | 2.76 (0.75), 0.016 |
| | L a-STS | 1.29, 0.272 | 0.86 (0.23), 0.492 |
| | L p-STS | 0.93, 0.359 | 0.69 (0.19), 0.492 |
| SE positive > Obj | L m-STS | 1.75, 0.291 | 1.95 (0.53), 0.223 |
| | L a-STG | 1.16, 0.334 | 1.44 (0.39), 0.265 |
| | L a-STS | 1.48, 0.291 | 1.30 (0.36), 0.265 |
| | L p-STS | -0.11, 0.912 | -0.60 (-0.17), 0.548 |
| SE negative > Obj | L m-STS | 3.06, 0.013 | 3.16 (0.86), 0.010 |
| | L a-STG | 1.76, 0.169 | 1.76 (0.48), 0.169 |
| | L a-STS | 0.98, 0.333 | 0.27 (0.07), 0.788 |
| | L p-STS | 1.20, 0.313 | 1.08 (0.29), 0.382 |

*Sensitivity analysis for spatial smoothing: each child's first-level model (including the valence-split model) was re-estimated on normalized but unsmoothed images with an otherwise identical design, and the group comparison was repeated on the 6 mm sphere values at the four Mellem et al. (2016) coordinates (two-sample t, df = 56; positive t indicates lower activity in the autistic group; d is Cohen's d; FDR p is corrected across the four regions within each contrast).*

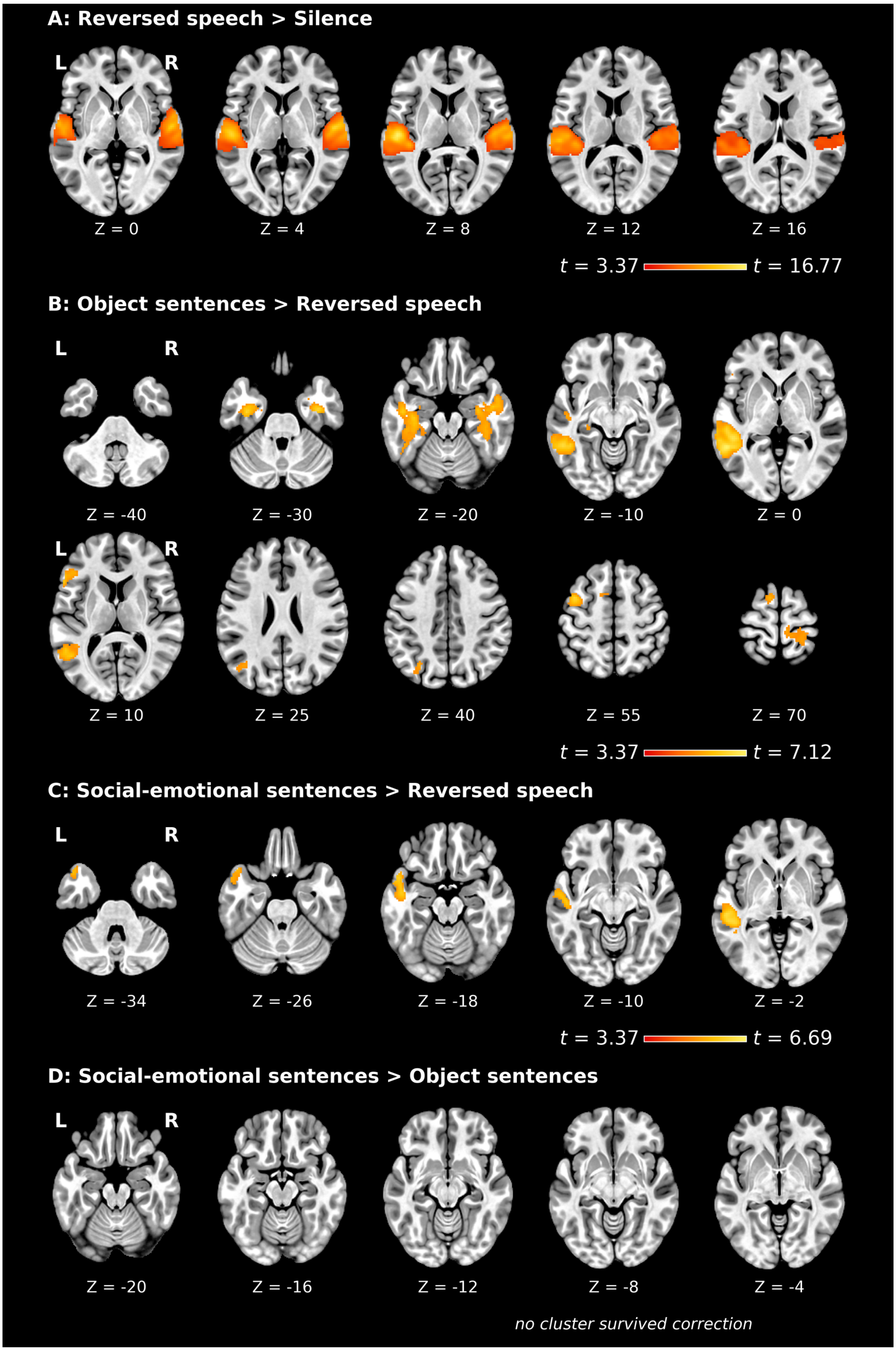


**Figure S1.** Whole-brain activity in autistic children ($n$ = 37). Second-level statistical maps for (A) Reversed speech > Silence, (B) Object sentences > Reversed speech, (C) Social-emotional sentences > Reversed speech, and (D) Social-emotional sentences > Object sentences, estimated in the autistic group. Clusters were formed at a voxel-level threshold of $p < 0.001$ ($t = 3.37$, $df = 32$) and are displayed only if they survived whole-brain cluster-level family-wise error correction at $p < 0.05$. No cluster survived correction for [Social-emotional > Object].

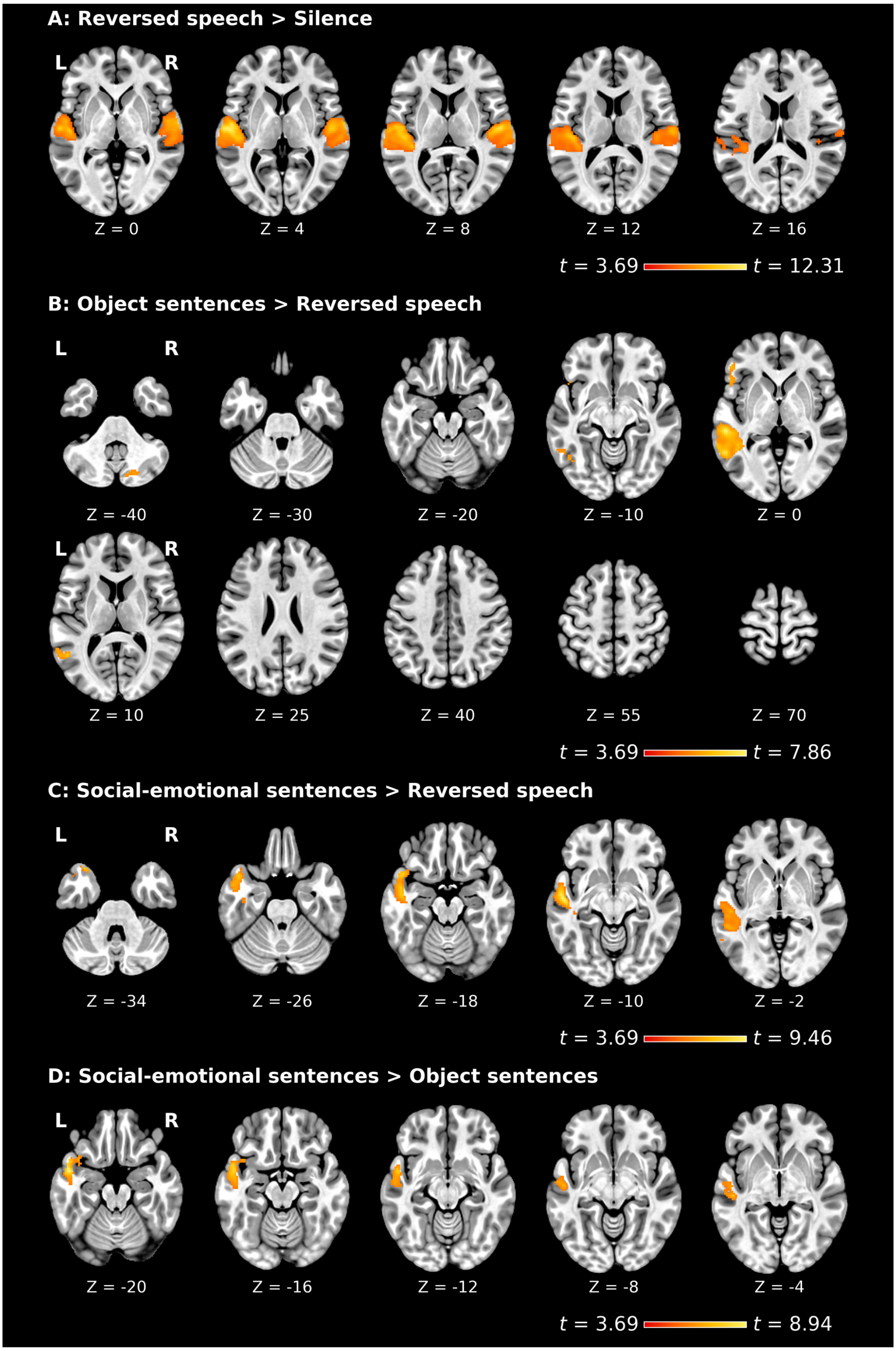


**Figure S2.** Whole-brain activity in non-autistic children ($n$ = 21). Second-level statistical maps for (A) Reversed speech > Silence, (B) Object sentences > Reversed speech, (C) Social-emotional sentences > Reversed speech, and (D) Social-emotional sentences > Object sentences, estimated in the non-autistic group. Clusters were formed at a voxel-level threshold of $p < 0.001$ ($t = 3.69$, $df = 16$) and are displayed only if they survived whole-brain cluster-level family-wise error correction at $p < 0.05$.

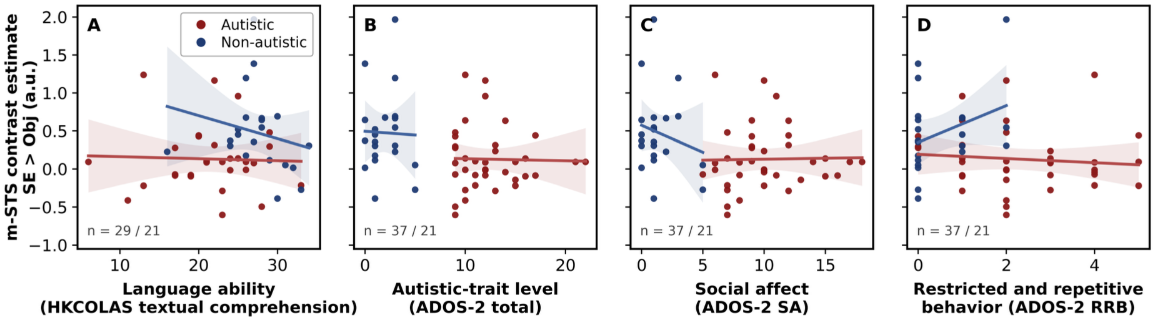


**Figure S3.** Left m-STS response to social-emotional over object sentences as a function of language ability and autistic-trait level. Each point is one child, colored by diagnostic group; lines are unadjusted within-group regressions with 95% confidence bands, drawn to show the raw data. HKCOLAS, Hong Kong Cantonese Oral Language Assessment Scale; ADOS-2, Autism Diagnostic Observation Schedule, Second Edition (Module 3); SA, social affect; RRB, restricted and repetitive behavior; SE, social-emotional sentences; Obj, object sentences; m-STS, middle superior temporal sulcus; L, left.
